# Combining dictionary- and rule-based approximate entity linking with tuned BioBERT

**DOI:** 10.1101/2021.11.09.467905

**Authors:** Ghadeer Mobasher, Lukrécia Mertová, Sucheta Ghosh, Olga Krebs, Bettina Heinlein, Wolfgang Müller

## Abstract

Chemical named entity recognition (NER) is a significant step for many downstream applications like entity linking for the chemical text-mining pipeline. However, the identification of chemical entities in a biomedical text is a challenging task due to the diverse morphology of chemical entities and the different types of chemical nomenclature. In this work, we describe our approach that was submitted for BioCreative version 7 challenge Track 2, focusing on the ‘Chemical Identification’ task for identifying chemical entities and entity linking, using MeSH. For this purpose, we have applied a two-stage approach as follows (a) usage of fine-tuned BioBERT for identification of chemical entities (b) semantic approximate search in MeSH and PubChem databases for entity linking. There was some friction between the two approaches, as our rule-based approach did not harmonise optimally with partially recognized words forwarded by the BERT component. For our future work, we aim to resolve the issue of the artefacts arising from BERT tokenizers and develop joint learning of chemical named entity recognition and entity linking using pre-trained transformer-based models and compare their performance with our preliminary approach. Next, we will improve the efficiency of our approximate search in reference databases during entity linking. This task is non-trivial as it entails determining similarity scores of large sets of trees with respect to a query tree. Ideally, this will enable flexible parametrization and rule selection for the entity linking search.

## I. Introduction

Name Entity Recognition (NER) is the first step in unlocking valuable information in unstructured and structured text data. Entity recognition is an essential step in many downstream applications such as entity linking, relation extraction, knowledge graph construction, etc. The main goal of chemical NER is to identify the boundaries of mentions of chemical entities from the text. It is challenging to build high-performance chemical NER tools due to the nature of the chemical text which contains abbreviations, Greek symbols, and various types of mathematical formulas (1).

Initially, NLP systems were mostly rule-based systems for knowledge extraction tasks. They have proven records since rule-based systems require less input data, however, the development of rule-based systems is challenging as it requires human intervention in the form of domain expertise to frame the rules. Moreover, it is very sensitive to small changes in the input data which makes it expensive and tedious. Machine learning systems brought some flexibility in developing NLP systems, as they learn rules during training and thereby overcame the laborious process of manual rule framing. Machine learning models still require feature engineering which again requires domain experts (34). This motivates the shift to Deep Learning (DL) models with the development of various DL models, such as Deep Neural Networks (DNN). DL architecture is flexible to be adapted to new problems and the same neural network can be applied to many various applications. However, DL requires a very large amount of data to outperform, causing expensive training due to complex data models (35).

In the conjunction of deep learning and rule-based approaches, hybrid approaches can leverage the frugality of expert rules for input data and the flexibility and generalizability of deep learning (36).

Most entity linking approaches disregard the entity recognition step, assuming that the correct annotations have been previously detected. The problem of Entity Linking in the chemical domain is strongly dependent on the rules of chemical nomenclature and habits in writing among the scientific community. This fact considerably complicates the automatic matching and searching of chemicals. There exist several tools to parse and convert a correctly formed systematic name into a standardized form, for example OPSIN (38). However, it does not consider any incompatibilities with IUPAC rules, typos, the generalization of chemically similar groups, the trivial names, and the existence of multiple meanings for one chemical name. The use of NLP techniques, on the other hand, lacks in handling domain-specific errors.

### A. Background

ChemNER (Chemical Named Entity Recognition) is presented as a classification task where the named entities in a given text are predicted. Chemical names are one of the in-demand entities to search in online search-engines like PubMed (14), which is used in varied application areas (17). Furthermore, the identification of the chemical entities continues to be a challenging task due to various reasons like non-standardized naming convention, naming ambiguities (5,7). So, there is a need for chemical databases for the efficient identification of ChemNER. As a result, currently, there are several gold-label chemical databases like BC5CDR (4), CHEMDNER (5), GENIA (15), CRAFT (8). There exist several silver-standard Chemical databases as well. In the case of the silver-standard corpus, the annotations achieved by multiple systems are then harmonized by establishing some rules in which a consensus is set for establishing (or not) a token as a certain entity class, using Deep Neural Networks (10). Although Giorgi & Bader (10) explored the area of BioNER, not ChemNER, with a comparable deep neural network, Habibi et al (12) performed ChemNER with other Biological NERs. However, they proposed a generic method based on deep learning and statistical word embeddings [Long Short-Term Memory network-Conditional Random Field (LSTM-CRF)], not a domain-specific word representation to Bio/ChemNER.

These word representations are also called word embeddings, such as Word2Vec (22,23) and GloVe(25). To represent a word, GloVe relies on the number of word co-occurrences over the given corpus whereas in Word2Vec the word representations are learned from a corpus in an unsupervised way without the word context, capturing the semantics of words throughout a two-layer neural network. Both representations are non-contextualized representations that can provide contextualized representation. Therefore, the state-of-the-art model BERT (Bidirectional Encoder Representations from Transformers) (9) is a contextualized word representation model that is based on a masked language model and pre-trained using bidirectional transformers (28). BioBERT(3) is a domain-specific language representation BERT model pre-trained on large-scale biomedical corpora. It adopts the same WordPiece tokenization to acquire the contextualized word representation. It is a subword-based tokenization algorithm that improves the coverage of the available dictionary (29). The performance of BioBERT for ChemNER reached F1=93.47, which was slightly better than that of the state-of-the-art PubMedBERT(11), which was F1=93.31. PubMedBERT explores the WordPiece tokenization to enrich the model with prefixes and suffixes of the ChemNERs, which played a crucial role to achieve comparable results to BioBERT. The precision of PubMedBERT model performance (94.26) is better than that of BioBERT (93.68). BioALBERT(32) outperforms all BERT-based models to recognize ChemNER. It adopted a self-supervised loss used in ALBERT (A Lite Bidirectional Encoder Representation from Transformers) that focuses on modelling inter-sentence coherence to better learn context-dependent representations. It incorporated parameter reduction techniques to lower memory consumption and increase the training speed in ChemNER. The F1 measure reaches 98.08 in this case, where the precision reaches 99.99.

In the community, the results are evaluated through the computation of precision, recall, and F1-measure. The evaluation is done through exact and approximate matching (24). Besides these machine learning-based models there exist rule-based methods as well.

Zhang & Elhadad (2013) developed an unsupervised solution that used three steps to address the NER task using TF-IDF scores, though the performances are not ashigh as in the case of a machine-learning-based model. The rule-based systems can perform with fewer data. Therefore, a hybrid approach can perform in a real-life scenario where sometimes domain-specific data is available, and sometimes not. Rocktäschel et al (2012) used a hybrid model with a merging result from two branches: one branch uses a CRF approach previously modelled (BANNER) aiming to focus on morphological complex structures such as IUPAC naming conventions. The other branch uses a ChemIDplus23(30) dictionary converted to a deterministic finite-state automaton for getting linear time complexity in text matching. There exist some pre-trained tools which can perform ChemNER. It is implemented as is the case of TaggerOne (18), tmChem(19). The state-of-the-art performance is found in the recent tool HunFlair(29).

Besides ChemNER recognition, the chemical normalization to a controlled vocabulary or an ontology is important for data integration for preparing the downstream tasks like relation extraction. The package pyMeSHSim (33) also recognizes ChemNEs by using MetaMap. It produces Unified Medical Language System (UMLS) concepts to map the UMLS concepts to Medical Subject Headings (MeSH) (38). It is a challenging task to achieve (13).

### B. Our Contribution

We perform joint learning of NER using fine-tuned BioBERT trained on chemical datasets and we perform EL using a rule-based system that assigns a unique (or none) MeSH ID to each Chemical Linking part, in order to leverage their relatedness and obtain a more generalizable system.

In this paper, we present our submitted, ongoing, work for BioCreative 7th edition for chemical identification task (2) “Track 2 - NLM-CHEM track Full-text Chemical Identification in PubMed articles.” The rest of the paper is organized as follows: first, we present our proposed two-stage approach in the Methods section. Then, we present our preliminary results, conclude, and discuss further work.

## II. Methods

This section focuses on the proposed methodology for BioCreative 7^th^ edition for track 2 focusing on identifying chemical entities and entity linking using MeSH. This section is composed of a two-stage approach (A) Chemical Identification (B) Chemical Entity Linking. Fig.1 depicts our proposed workflow.

**Fig. 1.**
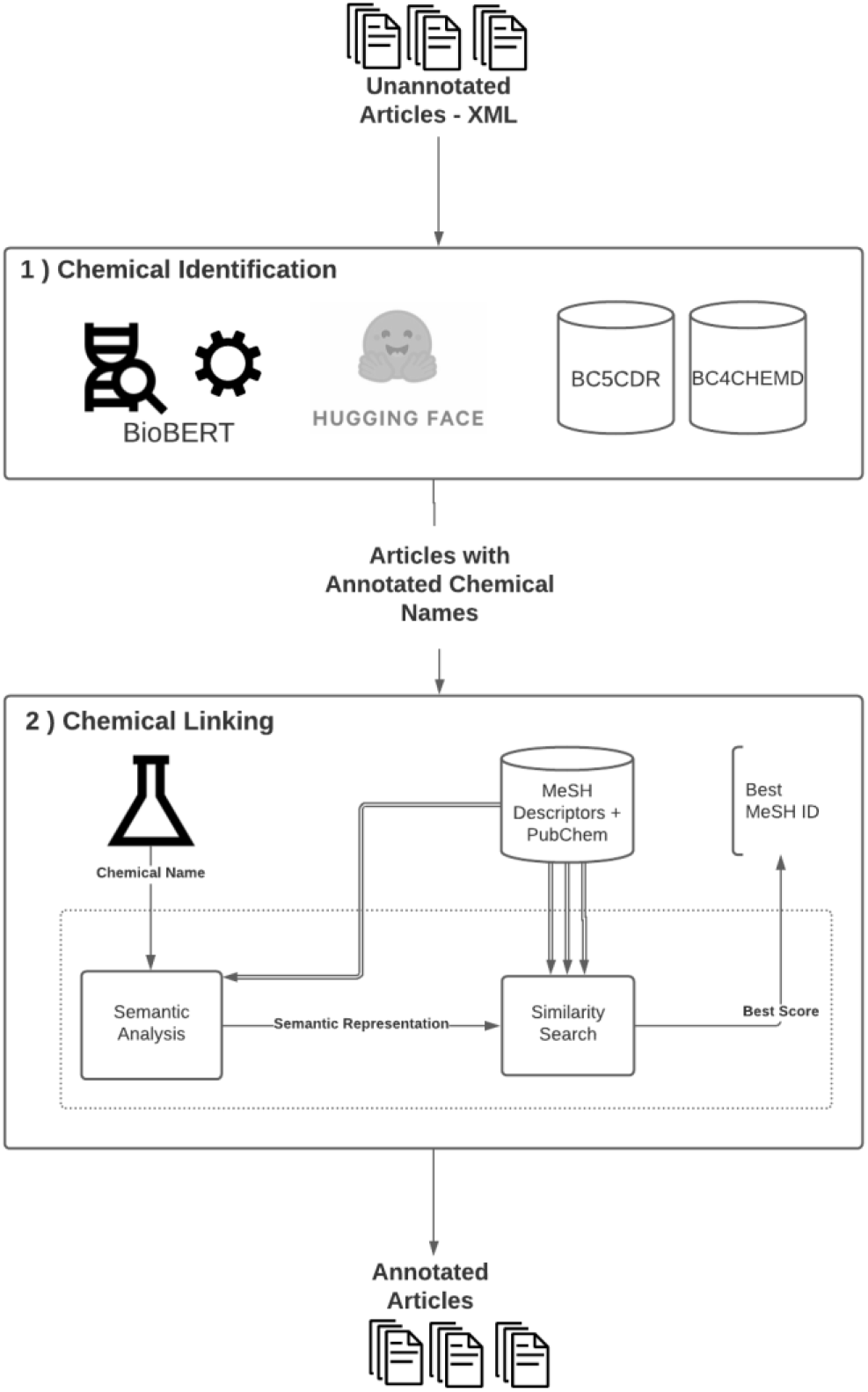
Proposed 2-staged approach focusing on the ‘Chemical Identification’ task. It consists of: (1) Chemical Identification, (2) Chemical Linking.

### A. Chemical Entity Identification

For identifying chemical entities, we have base ourselves on the most robust and first Biomedical pre-trained language model BioBERT (3). BioBERT uses self-supervised learning (SSL) performing mixed-Domain Pretraining (MDPT) on which the model was pre-trained over general domain text and then adapted to the biomedical domain.

Since this task mainly focuses on identifying chemical entities, we decided to use a fine-tuned BioBERT model enhanced by training using the BC5CDR-chemicals (4) and BC4CHEMD corpus (5). We conducted our work on one of HITS clusters “Cascade” on which we used one of its GPU nodes to run the model on the NLM-Chem corpora as well evaluating fine-tuned BioBERT using BC evaluation script.

Due to time limitations, we have used an existing model from Hugging Face 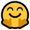 Hub developed by Alvaro Alonso and Carlos Badenes-Olmedo (6).

We have applied this fine-tuned BioBERT model on the chemical corpus of the NLM-Chem BioCreative VII track.

We conducted our work on a professional grade NVIDIA Tesla cluster, using one of its GPU nodes to run the model on the NLM-Chem corpora as well evaluating fine-tuned BioBERT using the BC evaluation script.

### B. Chemical Entity Linking

Our proposed solution for the Chemical Linking problem adapts a rule-based approach, developed by Mertová (37) to the problem at hand. The algorithm is flexible with respect to the input data. Its key concept is parsing the chemical entity into meaningful sub-word components. This tree is then standardized and compared to the trees corresponding to words in a reference database. These comparisons are approximate, allowing e.g. for spelling errors or errors of notation, producing a ranking of the reference database with respect to the chemical entity. We then picked the best match and applied a similarity cutoff.

The articles processed by the first module, Chemical Identification, represents the input to the Chemical Linking. Recognized chemicals are one by one processed by this workflow depicted in Figure 1, which shows the proposed workflow of the input name, the interconnection of the modules and communication with the reference database. Each chemical is analyzed and parsed via the Semantic analysis module. Its output is the complex tree structure containing semantic information about the chemical nature of the input chemical. Next, the parsed data structure is processed via a Similarity Search module which queries internally stored reference database.

#### i. Reference Database

A necessary part of the solution is the reference database preparation. For this purpose, we enriched the MeSH database of Descriptors (38) with synonyms from PubChem (39) database. Each MeSH descriptor in our resulting database contains its official name, the Entry Terms, and all available synonyms from PubChem. This composition of databases provides more accurate Entity Linking since the PubChem database consists of an immense amount of abbreviations and trivial names which are hard to derive automatically.

#### ii. Semantic Analysis

The Semantic Analysis module uses the knowledge and rules of organic chemistry, according to IUPAC. This knowledge allows chemical differentiating of each part of the compound names. Thus, we can separate individual components and assign different priorities according to their current position and type. It decomposes the input name into individual elements, each carrying specific structural and chemical information. Mertová (37) describes the algorithm in depth. The present work adapted the semantic analysis module, so it satisfied the nature of our input data. Also, we curated the internal lists of chemical constants, so the analysis provides a more precise match.

#### iii. Similarity Search

The considerable advantage of this Chemical Linking application is its acceptance of user-side errors. Similarity estimation is a computational process of evaluating the difference between two chemical compounds. The similarity estimation algorithm used in this module works with the structural and syntactic properties obtained from a chemical compound in the Analysis Module. The similarity estimation provided by the Similarity Search Module deals with the chemical-based errors and typos thanks to likening each chemical feature separately. It results in the similarity score value (difference) of two chemical names. This value is then processed in the last step, which is database searching. The searching result is a MESH entry with a minimal distance from our query.

## III. Results

In this section, we present our results for both chemical identification and linking as submitted to BioCreative Challenge 7^th^ edition. It is worth to mention, that the evaluation was done individually on each module separately.

### A. Chemical Entity Identification

We have evaluated the performance of the fine-tuned BioBERT which is trained on BC5CDR-chemicals and BC4CHEMD corpus. We have used the given evaluation script provided by BC Track 2. Table I. illustrates the performance of fine-tuned BioBERT on the NLM-Chem BioCreative VII track for strict and approximate evaluation respectively. However, when we investigated the numerous failure cases, we linked them to poor tokenization. BioBERT uses WordPiece tokenization. We spotted various issues related to out of vocabulary problems, meaningless sequences of sub-tokens and compound word problems.

**TABLE I.** Fine-tuned biobert model performance on BC7T2-NLMCHEM for strict and approximate evaluation

| Corpora | Precision | Recall | F1 | Time in seconds |
| --- | --- | --- | --- | --- |
| **Strict Evaluation** |  |  |  |  |
| BC7T2-NLMChem - Dev | 0.7210 | 0.6623 | 0.6904 | 0.247428 |
| BC7T2-NLMChem - Test | 0.7202 | 0.6322 | 0.6733 | 0.568769 |
| **Approximate Evaluation** |  |  |  |  |
| BC7T2-NLMChem - Dev | 0.8814 | 0.7915 | 0.8341 | 0.573672 |
| BC7T2-NLMChem - Test | 0.8669 | 0.7503 | 0.8044 | 1.300037 |

### B. Chemical Entity Linking

Our Chemical Linking approach can find many chemical names at the cost of complete accuracy. The accuracy of isolated, rule&dictionary based Chemical Linking on the data from the BC testing dataset was above 87%. The parameter set used remains untouched to the default setting. Due to time constraints we were unable to determine the compounded performance of the whole pipeline.

## IV. Conclusion and future work

In this paper, we presented a 2-stage approach that was tailored for the submission of Track 2 from the 7^th^ edition for BioCreative, reporting results for chemical identification and linking.

Although we used off-shelf fine-tuned BioBERT that is trained on chemical corpora, the quality of recognition is degraded. This mainly affects the new entities that are not part of the base vocabulary of BERT’s WordPiece tokenizer, resulting into multiple splitting of sub-tokens. We plan to investigate the possibility of resolving this drawback without increasing the size of the model. In addition, we need to extend our experimental settings to use ensembles of BioBERT and fine-tune them on NLM-chem corpora and other domain-specific corpora.

We plan to improve the accuracy of chemical linking in several ways. Firstly, we plan to fit some parameters, such as the acceptance threshold, to the training data. Furthermore, we plan to investigate the use of result presentations such as well-formed IUPAC structures that harmonize better with our rule-based approach. Better tuning will be possible with a faster search algorithm. This is currently our focus.

## Acknowledgement

Ghadeer Mobasher is part of the PoLiMeR-ITN (http://polimer-itn.eu/). Her project is supported by European Union’s Horizon 2020 research and innovation program under the Marie Skłodowska-Curie grant agreement PoLiMeR, No 81261. The work was supported by the Klaus Tschira Foundation, KTS. We thank the HITS IT services group for kind help. We would like to express our sincere gratitude to Robert Leaman and Rezarta Islamaj for their support and their encouragement to submit at this time. Finally, we would like to thank our colleagues “SDBVers” for their motivation and kind help.

